# Cell type specific long non-coding RNA targets identified by integrative analysis of single-cell and bulk colorectal cancer transcriptomes

**DOI:** 10.1101/2024.01.08.574618

**Authors:** Ante Mihaljevic, Philip Rubin, Panagiotis Chouvardas, Roberta Esposito

**Affiliations:** Department for BioMedical Research, University of Bern, 3008 Bern, Switzerland; Department of Medical Oncology, Inselspital, Bern University Hospital, University of Bern, 3010 Bern, Switzerland; Department for BioMedical Research, Urology Research Laboratory, University of Bern, 3008 Bern, Switzerland

## Abstract

Long non-coding RNAs (lncRNAs) represent an emerging class of genes which play significant and diverse roles in human cancers. Nevertheless, the functional repertoires of lncRNAs in cancer cell subtypes remains unknown since most studies are focused on protein coding genes. Here, we explored the contribution of lncRNAs in Colorectal Cancer (CRC) heterogeneity. We analyzed 49,436 single-cells from 29 CRC patients and showed that lncRNAs are significantly more cell type specific compared to protein-coding genes. We identified 996 lncRNAs strongly enriched in epithelial cells. Among these, 98 were found to be differentially expressed in tumor samples compared to normal controls, when integrating 270 bulk CRC profiles. We validated the upregulation of two of them (CASC19 and LINC00460) in CRC cell lines and showed their involvement in CRC proliferation by CRISPR-Cas9 knock down experiments. This study highlights a list of novel RNA targets for potential CRC therapeutics, substantiated through experimental validation.

## Introduction

Colorectal cancer (CRC) is the third most common cancer and the second leading cause of cancer-related deaths worldwide^1^. Tumor heterogeneity, a hallmark of CRC, has emerged as a critical factor complicating effective treatment strategies. Within a single tumour mass, CRC exhibits remarkable cellular and molecular diversity, with distinct subpopulations of cancer cells exhibiting varied phenotypes, genotypes, and functional properties^2^. Intratumoral heterogeneity complicates accurate diagnosis and prognosis, as well as significantly impacts treatment response and resistance mechanisms^3,4^. To gain deeper insights into the complex landscape of CRC, researchers have turned to emerging technologies such as single-cell RNA sequencing (scRNA-seq). By enabling transcriptomic profiling of individual cells within a tumor, scRNA-seq represents a powerful approach to interrogate the heterogeneity present within the tumor microenvironment^5^. This tool provides a comprehensive overview of the diverse cell types present in a tumor sample as well as their functional states, allowing for the identification of key drivers and potential therapeutic targets.

A rich mine of new potential CRC targets can be explored in the gene class of long non-coding RNAs (lncRNAs)^6^. These non-coding RNA molecules, once thought to be transcriptional noise, have now emerged as key regulators of gene expression and cellular processes^7^. Dysregulation of lncRNAs has been implicated in many cancers including CRC, affecting various pathophysiological processes such as tumor initiation, progression, and metastasis^8^. For instance, CCAT1 (colon cancer-associated transcript 1) has been identified as an oncogenic lncRNA that promotes CRC growth and metastasis through multiple mechanisms, including chromosomal looping and gene regulation^9^. BANCR (BRAF-activated non-coding RNA) is another lncRNA that has been implicated in CRC progression and metastasis, acting as a sponge for microRNAs and modulating the expression of target genes^10,11^. Due to their cell type-specific expression and function^6,12^, LncRNAs present promising candidates for targeted therapies and precision medicine approaches. To date, little is known about lncRNA expression in the intra-tumor heterogeneity.

Here, through integrating cutting-edge approaches, we delve into the intricate relationship between tumor heterogeneity and lncRNAs in CRC. We analyzed available scRNA-seq data, providing the panorama of epithelial lncRNAs. Up-regulated epithelial lncRNAs in CRC compared to normal tissues were identified by using bulk RNA-sequencing CRC profiles. Ultimately, we used CRISPRi technology to validate the involvement of two lncRNAs, CASC19 and LINC00460, in CRC disease progression, thereby demonstrating the potential of our approach in the identification of functional lncRNAs and novel RNA drug targets.

## Results

### LncRNAs are a more specific cell type marker compared to protein coding genes

Aiming to explore the role of lncRNAs in CRC heterogeneity, we analyzed single-cell RNA sequencing (scRNA-seq) data from 29 samples, derived from tumor, border regions or matched normal mucosa. After quality control we analyzed 49’436 single-cell transcriptomes, measuring 28’751 genes. The single-cell profiles are of high quality as is reflected by the number of features and low mitochondrial content (**Suppl. Fig. 1A)**. Uniform Manifold Approximation and Projection (UMAP) analysis shows that the cells cluster by cell type (**Fig. 1A**), as defined in the original publication. No batch effect was observed among the two datasets after merging them (**Suppl. Fig. 1B**). The cells were re-annotated by an automated method and the cell type labels were checked for consistency between the two approaches (**Suppl. Fig. 1C**). Hence, we concluded that this is a high-quality dataset, with no batch effect and with accurate cell type annotation labels. We then sought to explore the cell type specific expression of protein-coding and lncRNA genes by estimating the Gini coefficient of their expression in the different cellular compartments. LncRNAs show a strikingly higher Gini coefficient of expression compared to protein-coding genes (**Fig. 1B**), suggesting that they are overall expressed in a much more cell type specific manner. As an example, LINC00523 has a Gini coefficient of 1, suggesting that it is solely expressed in only one cell type. Indeed, when estimating the density of expression of LINC00523, it is only found in epithelial cells (**Fig. 1C**). Differential expression analysis was then performed in order to identify lncRNAs enriched in epithelial cells. We identified 996 lncRNAs that show a clear overexpression in epithelial cells while they are only mildly expressed in the other cell types (**Fig. 1D**), generating a unique list of putative RNA targets enriched in CRC epithelium.

**Figure 1.**
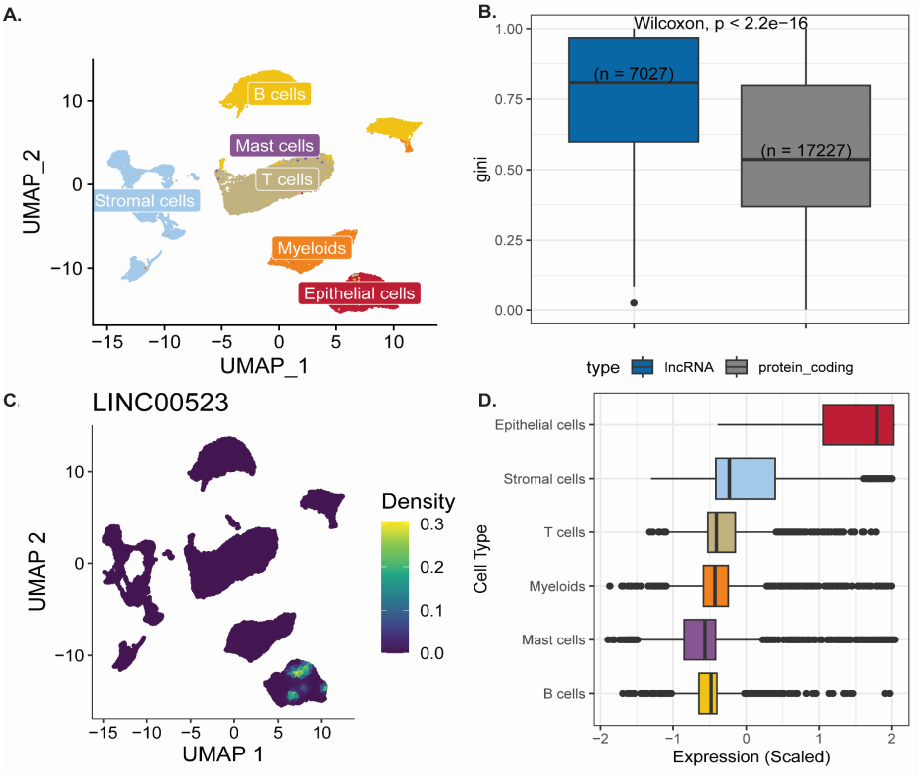
ScRNA-seq and cell type enrichment analysis. A. UMAP of the single cells colored by cell type. B. Comparing cell type specificity between lncRNAs and protein coding genes. C. Example of a lncRNA with high Gini coefficient. D. Expression overview of epithelial lncRNAs across cell lines.

### Integration of scRNA-seq data with large-scale transcriptomics datasets reveals disease relevant lncRNAs

To further prioritize the list of epithelial lncRNAs and identify disease relevant genes, we accessed differential gene expression results from The Cancer Genome Atlas (TCGA) consortium, based on 270 bulk RNA-sequencing profiles. We identified 426 differentially expressed lncRNAs, from which 303 are down- and 126 are up-regulated in the tumor samples. When we overlap these genes with the identified 996 epithelial lncRNAs, we observe 98 common genes (77 up- and 21 down-regulated) (**Fig. 2A/B**). We hypothesize that promising drug targets should show a minimal expression in normal tissue to further reduce the probability of unintended side effects.

**Figure 2.**
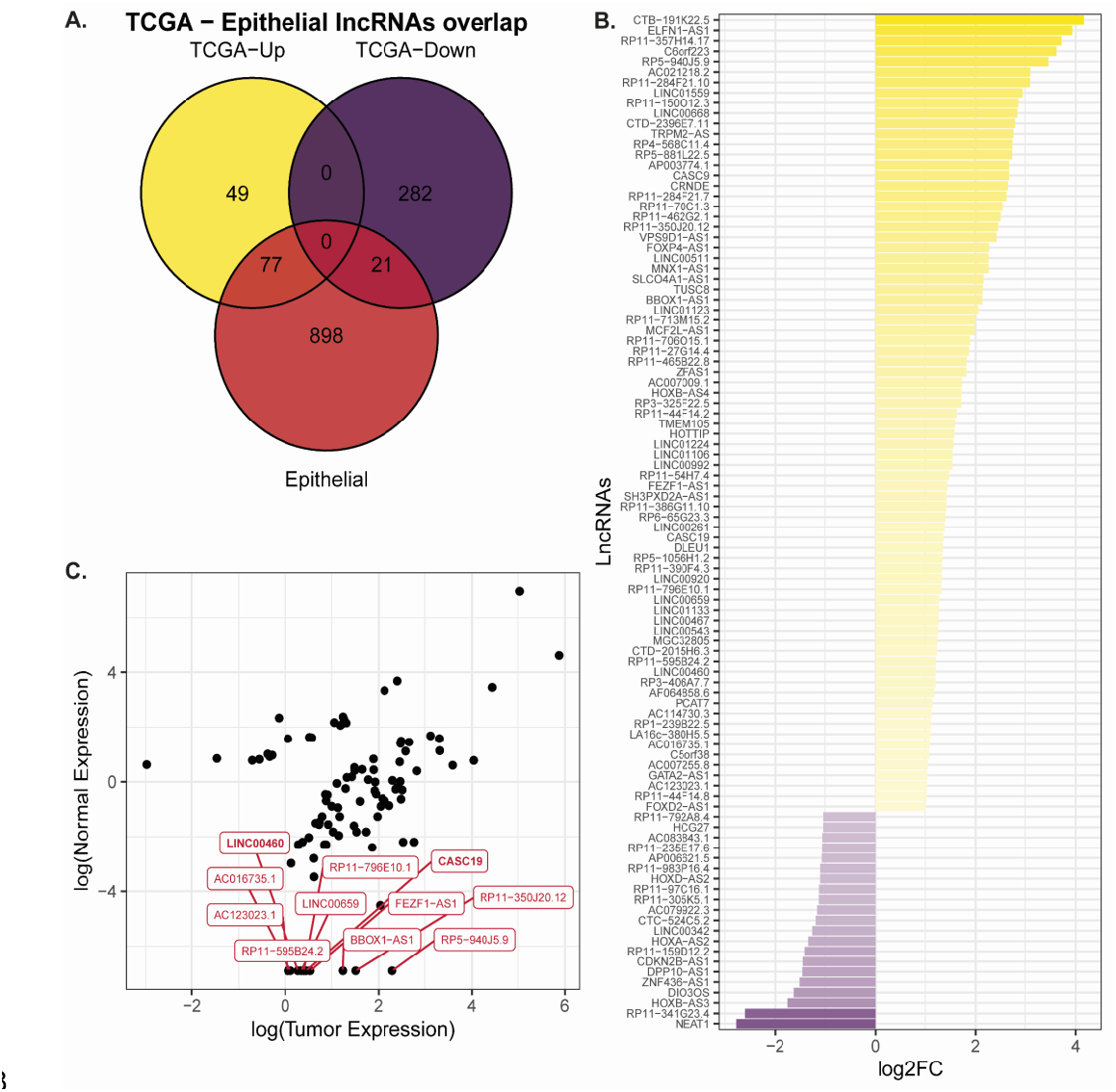
Integration with TCGA and prioritization of lncRNAs. A. Overlap of epithelial (red) and differentially expressed lncRNAs in TCGA (yellow/purple). B. TCGA differential expression analysis of overlapping lncRNAs. C. Tumor vs normal expression highlighting 11 lncRNAs which show minimal expression in normal tissues.

Therefore, from the 77 epithelial lncRNAs which were upregulated in tumor samples we prioritized 11 candidates: AC016735.1, AC123023.1, BBOX1−AS1, CASC19, LINC00659, FEZF1−AS1, LINC00460, RP11−595B24.2, RP11−796E10.1, RP5−940J5.9 and RP11−350J20.12 (**Fig. 2C**). Taken together, these lncRNAs are enriched in the epithelial cells of CRC patients, upregulated in the tumor and minimally expressed in normal tissue. Therefore, they represent highly promising candidates as novel CRC RNA drug targets.

### Validating the expression of prioritized lncRNAs in CRC cell lines and its efficient reduction through CRISPR knockdown

To ensure the reliability of our results, we selected two lncRNAs from the 11 that show a specific expression in CRC samples (**Fig. 2C**) for further validation: *CASC19* (Cancer Susceptibility candidate 19; ENSG00000254166.2) and *LINC00460* (long intergenic non-protein coding RNA 460; ENSG00000233532.5).

CASC19 is a lncRNA that is located on chromosome 8q24.2, containing at least 29 different annotated transcripts in Gencode v38 annotation^13^ (**Fig. 3A**). CASC19 has been described as an oncogenic gene involved in CRC progression^14,15^. In normal tissues, CASC19 is expressed mainly in esophagus (**Suppl. Fig. 2A**), while in cancerous tissues, especially in CRC, it is upregulated (**Suppl. Fig. 2B**).

**Figure 3.**
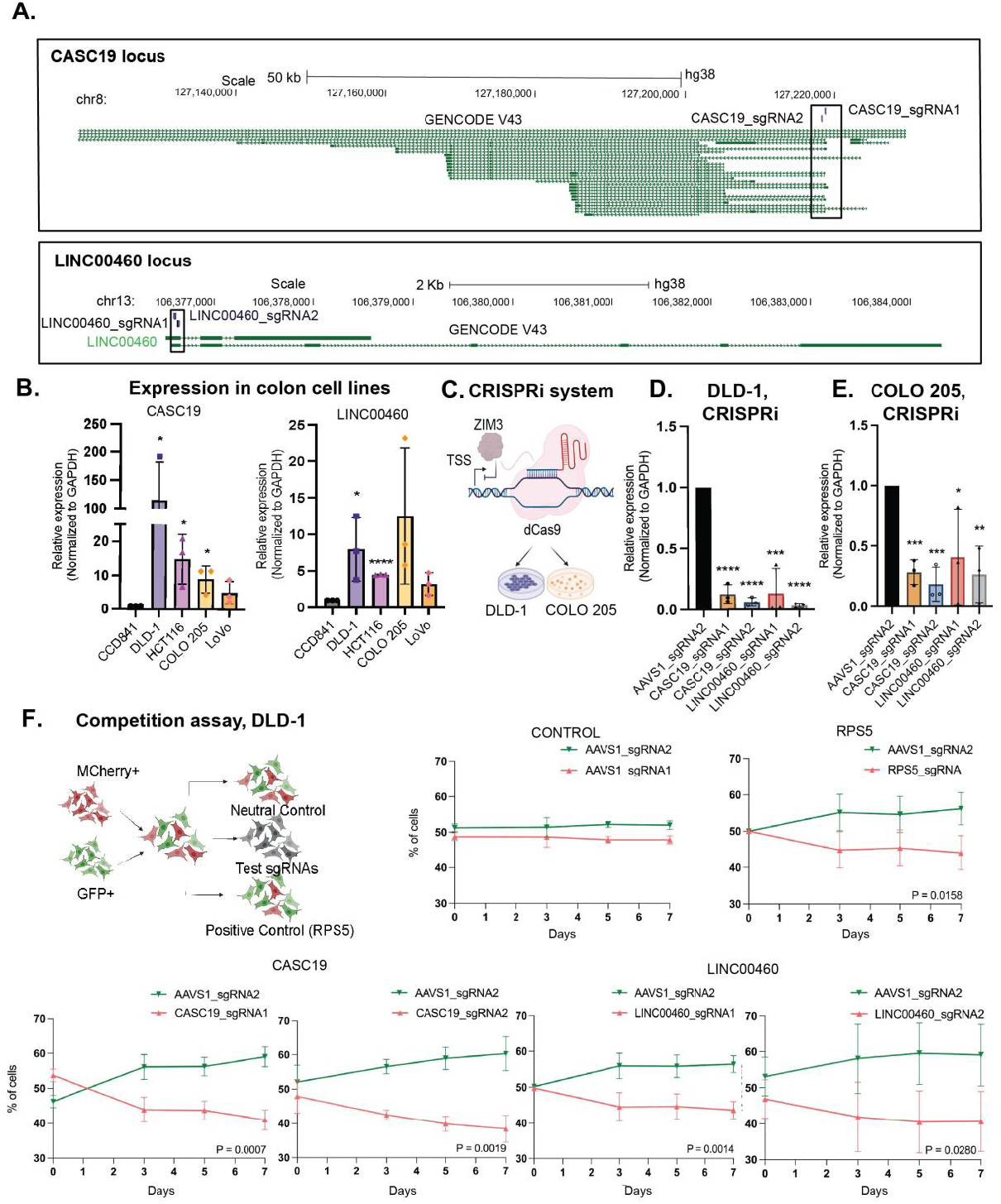
Expression of CASC19 and LINC00460 in CRC cell lines and their involvement in CRC proliferation. A. The genomic locus of CASC19 and LINC00460 lncRNA genes. The sgRNA position is highlighted by the black boxes. B. The expression levels of CASC19 and LINC00460 assessed in various CRC cell lines relative to the normal epithelial colon cell line CCD841. C. Cartoon of CRISPRi system consisting of dCas9 enzyme fused with ZIM3-derived KRAB repressor domain. D. and E. Expression levels of CASC19 and LINC00460 genes in DLD-1 and COLO 205 cell lines after CRISPRi knockdown. Each gene was targeted using two distinct sgRNAs. F. Competition assay used to assess the phenotypic implications of CASC19 and LINC00460 CRISPRi downregulation. Initially, cells expressing the mCherry marker and carrying various single-guide RNAs (sgRNAs) were combined with GFP-positive control cells in equal proportions. The red-to-green fluorescence ratio was measured at consecutive time intervals using flow cytometry. Data are plotted as mean ± SD (n=3). Statistical significance was estimated by two-tailed (B) or one-tailed (D, E) Student’s t test (*p < 0.05, **p < 0.01, ***p < 0.001, ****p < 0.0001).

LINC00460 is an intergenic lncRNA, located on chromosome 13q33.2, which has been previously described in human malignancies^16^. It contains two annotated transcripts, partially sharing the first exon: a shorter one containing three exons made up of 1’746 nucleotides and a longer one of seven exons made up of 2’124 nucleotides base pairs, mainly expressed in normal brain tissue (**Suppl. Fig. 2C**). They both show a significant upregulation in CRC tissues when compared to healthy tissues (**Suppl. Fig. 2D**).

We assessed the expression levels of CASC19 and LINC00460 genes in a pool of colon cell lines: DLD-1, COLO 205, HCT116, and LoVo as colon cancer models and CCD841 as a normal epithelial cell line. Both genes are upregulated in CRC cell lines compared to the normal epithelial colon cell line CCD841 with the expression being the most pronounced in DLD-1 cell line (on average 114.57 fold change for CASC19 and 7.98 fold change for LINC00460) (**Fig. 3B)**.

### Competition assay indicates CASC19 and LINC00460 are involved in CRC proliferation

To assess the impact of these genes on cancer cell phenotype, we proceeded with a CRISPRi-mediated knockdown (**Fig. 3C**) in the DLD-1 and COLO205 cell lines. We established DLD-1 and COLO205 to stably express ZIM3-KRAB-BFP-dCas9 (enriched for BFP expression; **Suppl. Fig. 3A**). The KRAB domain derived from the ZIM3 protein was selected as a repressor domain based on its notable capability to induce robust knockdown effects without causing non-specific impacts on cell viability and gene expression^17^. We transduced these cell lines with lentiviral vectors carrying different sgRNAs. Each gene was targeted with two different high-scoring sgRNAs, that mediated a potent downregulation of CASC19 and LINC00460 genes in both cell lines (**Fig. 3D/E**). However, a more pronounced reduction in gene expression was observed in the DLD-1 cell line **(Fig. 3D)** which was then used to assess the impact of gene downregulation on growth phenotype.

Growth phenotypes resulting from CRISPRi-mediated knockdown of *CASC19* and *LINC00460* genes were tested by competition assay. Cells transduced with sgRNAs targeting both lncRNAs were tagged with mCherry, while cells transduced with control were tagged with GFP (**Fig. 3F, Suppl. Fig. 3B**). mCherry+ and GPF+ cells are mixed in a 1:1 proportion on day 0 and the change in ratio of mCherry+ to GPF+ cells was followed over a one-week period with measurements taken at two-day intervals. We could show that both sgRNAs for CASC19 and LINC00460 yielded strong effects on cell fitness: mCherry+ cells expressing sgRNAs targeting our candidates were out-competed by control cells (GFP+, targeting AAVS1 locus). The effect was comparable to inhibition of the essential ribosomal gene RPS5, highlighting the bona fide of our approach to identify valuable lncRNA targets in epithelial cancer cells.

## Discussion

Despite the some advantages, most scRNA-seq studies were still limited to a focus on protein-coding genes and overlook the potential role of lncRNAs. This study aimed to investigate the role of long non-coding RNAs (lncRNAs) which have emerged as central players and key regulators in multiple biological processes such as cancer cell proliferation and drug resistance^8,18^ in colorectal cancer (CRC) heterogeneity.

By analyzing a large dataset of single-cell and bulk transcriptomics profiles, we demonstrated the cell type-and tissue-specificity of lncRNAs in CRC and identified the first set of lncRNAs enriched in CRC epithelial cells. Targeting such cell-type enriched lncRNAs could potentially offer novel therapeutic strategies that exploit the unique vulnerabilities of CRC epithelial cells. Moreover, we focused on the differential expression of certain lncRNAs in tumor versus normal samples and validated their functional involvement in CRC proliferation.

Our work serves as a foundation for future investigations focusing on the functional characterization of these epithelial-enriched lncRNAs and their potential implications in CRC progression. This study highlights the importance of considering lncRNAs in understanding the heterogeneity and functional diversity of CRC cells.

Moreover, we focused on two lncRNAs, CASC19 and LINC00460, which showed upregulation in CRC cell lines. Both CASC19 and LINC00460 are known to exhibit upregulation in CRC tissues, particularly in samples from patients with aggressive forms of CRC, thereby they are linked with poorer patient prognosis^14,15^. CASC19 is involved in enhancing the invasive and migratory potential of CRC cells through positive regulation of cell migration inducing hyaluronidase 1 (CEMIP) and epithelial-mesenchymal transition markers^19^. Similarly, LINC00460 is involved in modulating cell migration and invasion by regulating miR‐ 613/SphK1^20^ and miR-939-5p/LIMK2 axes^21^. Through CRISPR knockdown experiments, we substantiated experimental evidence for their functional involvement in CRC proliferation in multiple cell models. These findings suggest that CASC19 and LINC00460 may serve as promising therapeutic targets for CRC treatment. Further investigation is warranted to elucidate their precise molecular mechanisms and evaluate their potential as druggable targets.

Furthermore, we propose the integration of bulk and single-cell transcriptomics datasets to achieve a more comprehensive understanding of CRC heterogeneity. This integration approach addresses the limitations of solely relying on either bulk or single-cell data, providing a more robust and comprehensive view of CRC transcriptomic profiles. Single-cell analysis provided valuable insights into the cellular composition and heterogeneity of CRC, while bulk RNA-sequencing allowed us to explore differential expression patterns across a larger cohort of patients.

While this study provides important insights into the role of lncRNAs in CRC heterogeneity and highlights potential therapeutic targets, there are limitations to consider. Firstly, lncRNAs’ low overall expression and incomplete annotation catalogues could potentially hinder their prioritization in scRNA-seq datasets. In future studies, this limitation can be addressed by performing *de novo* transcriptome assembly in matched bulk transcriptomes as was shown in other contexts^22^. Moreover, the functional experiments are focused on a subset of identified lncRNAs, and further studies are needed to explore the functional significance of other differentially expressed lncRNAs. Finally, experimental validation in a cohort of patient samples would strengthen the findings, possibly utilizing more clinically relevant experimental models such as Patient Derived Organoids (PDOs).

In conclusion, this study sheds light on the cell-type specificity of lncRNAs in CRC, revealing their potential as diagnostic markers and therapeutic targets. The identified lncRNAs enriched in epithelial cells and their experimental validation highlight the functional relevance of lncRNAs in CRC proliferation. The integration of single-cell and bulk transcriptomics datasets provides a comprehensive view of CRC heterogeneity and offers a data-driven approach to identify cell-type enriched RNA drug targets. Further investigation and validation of these targets may open new avenues for precision medicine approaches in the treatment of CRC.

## Materials and Methods

### Single-cell and bulk RNA sequencing data and analysis

Single-cell RNA sequencing data were downloaded from Gene Expression Omnibus, with accession numbers GSE132465 and GSE144735^23^. Raw data were pre-processed keeping those cells which express at least one gene and have less than 5% mitochondrial reads and those genes which are expressed in at least one cell. The two datasets were merged resulting in 49’436 single cells and 28’751 measured genes. Downstream single-cell analysis was performed using Seurat^24^ (version 4.1.1). Cell type annotation was assessed by the original publication and validated by scMRMA^25^ (version 1.0). Gene types were assessed by parsing the Gencode annotation^13^ (version 38). Cell type specificity was estimated by calculating the Gini coefficient via the DescTools package (version 0.99.46). TCGA data were assessed via GEPIA2^26^ web server. All analyses were performed in R (version 4.1.2). Visualizations were generated using the R packages ggplot2 (version 3.4.0), pals (version 1.7) and ggvenn (version 0.1.9). The code for the analysis and generation of the figures can be found in github https://github.com/pchouvardas/COAD_lncRNAs.

### Cell line maintenance

DLD-1 and COLO-205 were purchased at ATCC and regularly tested for mycoplasma infection. Cells were grown in RPMI-1640 Medium (Sigma) supplemented with 10% (v/v) Fetal Bovine Serum (Gibco), 2mM L-glutamine (Thermo Fisher), 100 Units/mL penicillin (Thermo Fisher) and 100 mg/mL streptomycin (Thermo Fisher). HCT116 cells were grown in McCoy’s 5A Medium (Sigma) supplemented with 10% (v/v) Fetal Bovine Serum (Gibco), 2mM L-glutamine (Thermo Fisher), 100 Units/mL penicillin (Thermo Fisher) and 100 mg/mL streptomycin (Thermo Fisher). LoVo cells were grown in Kaighn’s Modification of Ham’s F-12 Medium (Sigma) supplemented with 10% (v/v) Fetal Bovine Serum (Gibco), 2mM L-glutamine (Thermo Fisher), 100 Units/mL penicillin (Thermo Fisher) and 100 mg/mL streptomycin (Thermo Fisher). The cells were grown at 37°C in the presence of 5% CO_2_. HEK293T cells and CCD841 cells were grown in Dulbecco’s Modified Eagle’s Medium (Sigma) supplemented with 10% (v/v) Fetal Bovine Serum (Gibco) and 2mM L-glutamine (Thermo Fisher) 100 Units/mL penicillin (Thermo Fisher) and 100 mg/mL streptomycin (Thermo Fisher. All the cells were grown at 37°C in the presence of 5% CO_2_.

### Lentivirus production

24 hours before transfection 2.5 x 10^6^ HEK293T cells were seeded 100 mm tissue culture-treated plates coated with 0.1 mg/L poly-D-lysine (Gibco). Cells were triple-transfected with 12.5 μg of the plasmid of interest (UCOE-SFFV-ZIM3-KRAB-dCas9-P2A-BFP plasmid or pDECKO_mCherry plasmid containing sgRNA), 4 μg of pCMV-VSV-G (Addgene #8454) and 7.5 μg of psPAX2 (Addgene #12260) which were mixed with Lipofectamine 2000 (Thermo Fisher) and Opti-MEM™ I Reduced-Serum Medium (Gibco) according to the manufacturer instructions. 4-6 hours after transfection, the medium was replaced with complete DMEM (Gibco). Viral supernatant was collected 24-, 48- and 72-hours post-transfection. Collected viral supernatants were eventually pooled together and centrifuged at 3000 rpm for 15 min. The supernatant was collected and mixed with cold PEG-it Virus Precipitation Solution (4x). The mix was refrigerated overnight at 4°C and centrifuged at 1500 g for 30 min at 4°C. The supernatant was removed, followed by another round of centrifugation at 1500g for 5 min. Any remaining supernatant was removed, and the lentiviral pellet was suspended in 200μL of cold 1x PBS (Gibco), aliquoted into cryogenic vials and stored at -80°C.

### Generation of cell lines stably expressing dCas9

DLD-1 and COLO-205 cell lines stably expressing dCas9 fused to ZIM3 repressor domain, were generated through transduction of parental DLD-1 and COLO 205 cells (1.5 x 10^6^) with a concentrated lentiviral vector (30μL) carrying UCOE-SFFV-Zim3-dCas9-P2A-BFP plasmid suspended in 10mL of complete RPMI-1640 medium (Gibco) containing 8 μg/mL Polybrene. BFP-positive cells were selected through several rounds of expansion and FACS sorting.

### Design and cloning of individual sgRNAs

sgRNAs for CRISPRi were designed using the CRISPick tool from Broad Institute (https://portals.broadinstitute.org/gppx/crispick/public). For each sgRNA, forward and reverse DNA oligos were synthesised with 20bp overlaps with the 3’ end of the U6 promoter and the 5’ end of the sgRNA scaffold sequence. For cloning of annealed oligos into the pDecko_mCherry backbone (Addgene #78534), the plasmid was first digested with BsmBI, loaded on a 0.8% agarose gel and purified. 100ng of plasmid was combined with 1mL of each DNA oligo (1mM), and 10mL of homemade Gibson Assembly Mastermix (2x). The cloning product was checked by Sanger sequencing. Validated plasmid constructs were transformed into competent Stbl3™ *E. coli* cells - prepared with Mix & Go! *E. coli* Transformation Kit (Zymo Research). Plasmids were prepared following standard protocols. Generated plasmids were then used for lentivirus production.

sgRNA sequences are listed in Supplementary Table 1.

### Knockdown of lncRNA genes in CRC cell lines stably expressing dCas9

DLD-1 and COLO 205 cell lines stably expressing sgRNA targeting LINC00460 and CASC19, lncRNA genes were generated through transduction of parental DLD-1 and COLO 205 cells (0.5 x 10^6^) with a concentrated lentiviral vector (10μL) carrying pDecko_mCherry suspended in 2mL of complete RPMI-1640 medium (Gibco) containing 8 μg/mL Polybrene. 24 hours post-transduction, the antibiotic selection was induced by supplementing the culturing medium with 2 mg/mL puromycin (Thermo Fisher) for at least 3 days.

### RT-qPCR gene expression analysis

Total RNA was extracted from 500’000 cells of each cell line with Quick-RNA™ Miniprep Kit (Zymo Research). Isolated RNA was then retro-transcribed into cDNA with GoScript™ Reverse Transcription System (Promega) and the expression of the CRISPRi targeted lncRNA genes was assessed through Real-Time PCR with the GoTaq® qPCR Master Mix (Promega). Primers sequences are listed in Supplementary Table 1.

### Competition assay

After transduction with lentiviral particles containing pDecko_mCherry sgRNA constructs targeting LINC00460 and CASC19 and seven days after Puromycin selection (2 μg/mL), 250 000 cells stably expressing effector and sgRNA were mixed with 250 000 GFP expressing cells. Cell growth was evaluated by measuring the ratio of GFP- and mCherry-positive cells over seven days by flow cytometry on LSR II SORP instrument (BD Biosciences). As a positive control, cells were established to stably express the CRISPRi effector in conjunction with a single guide RNA (sgRNA) that specifically targeted the essential ribosomal gene RPS5. Conversely, as a negative control, cells were also established to stably express the CRISPRi effector along with an sgRNA that targeted the AAVS1 locus.

## Acknowledgements

The results of the bulk RNA-sequencing data utilized in this study were based upon data generated by the TCGA Research Network: https://www.cancer.gov/tcga.

The plasmid UCOE-SFFV-Zim3-dCas9-P2A-BFP (Addgene #188767) was a kind gift of Marco Jost’s lab.

We would like to thank Beatrice-Ederer Weber and Helmut-Horten foundations for the financial support. Additionally, we extend our thanks to the FACS facility of the University of Bern for their technical expertise.

## Author contributions

A.M. and P.R. performed the wet lab experiments. P.C. performed bioinformatic analysis. A.M. and R.E. designed and monitored experiments. All authors analyzed and interpreted data. P.C. and R.E. wrote the manuscript and all authors contributed to the review and editing of the manuscript.

## Data availability statement

Single-cell RNA sequencing data were downloaded from Gene Expression Omnibus, with accession numbers GSE132465 and GSE144735^23^. TCGA data were assessed via GEPIA2^26^ web server.

All the data generated in this study can be found https://github.com/pchouvardas/COAD_lncRNAs.

## Code availability statement

The code for the analysis and generation of the figures can be found in github https://github.com/pchouvardas/COAD_lncRNAs.

