## Supplementary Figures for "Cell type specific long non-coding RNA targets identified by integrative analysis of single-cell and bulk colorectal cancer transcriptomes"

**Suppl. Figure 1**

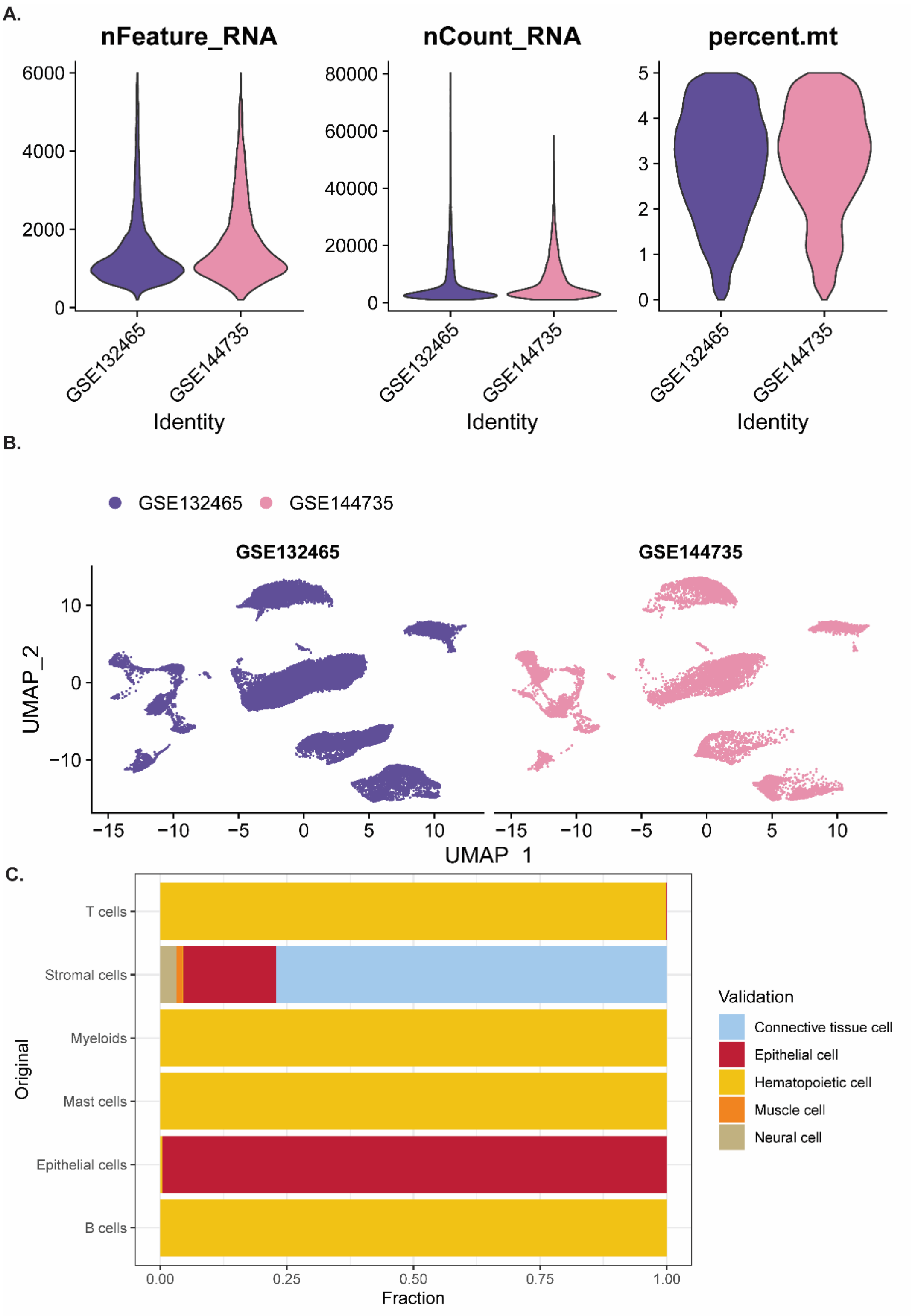


**Suppl. Figure 1. ScRNA-seq and cell type enrichment analysis.** A. Pre-processing descriptive statistics of the scRNA-seq datasets; number of features, number of counts and percent of mitochondrial reads. B. UMAP split by dataset shows no apparent batch effect. C. Overlap of cell type annotations compared to novel ones generated by using scMRMA.

**Suppl. Figure 2**

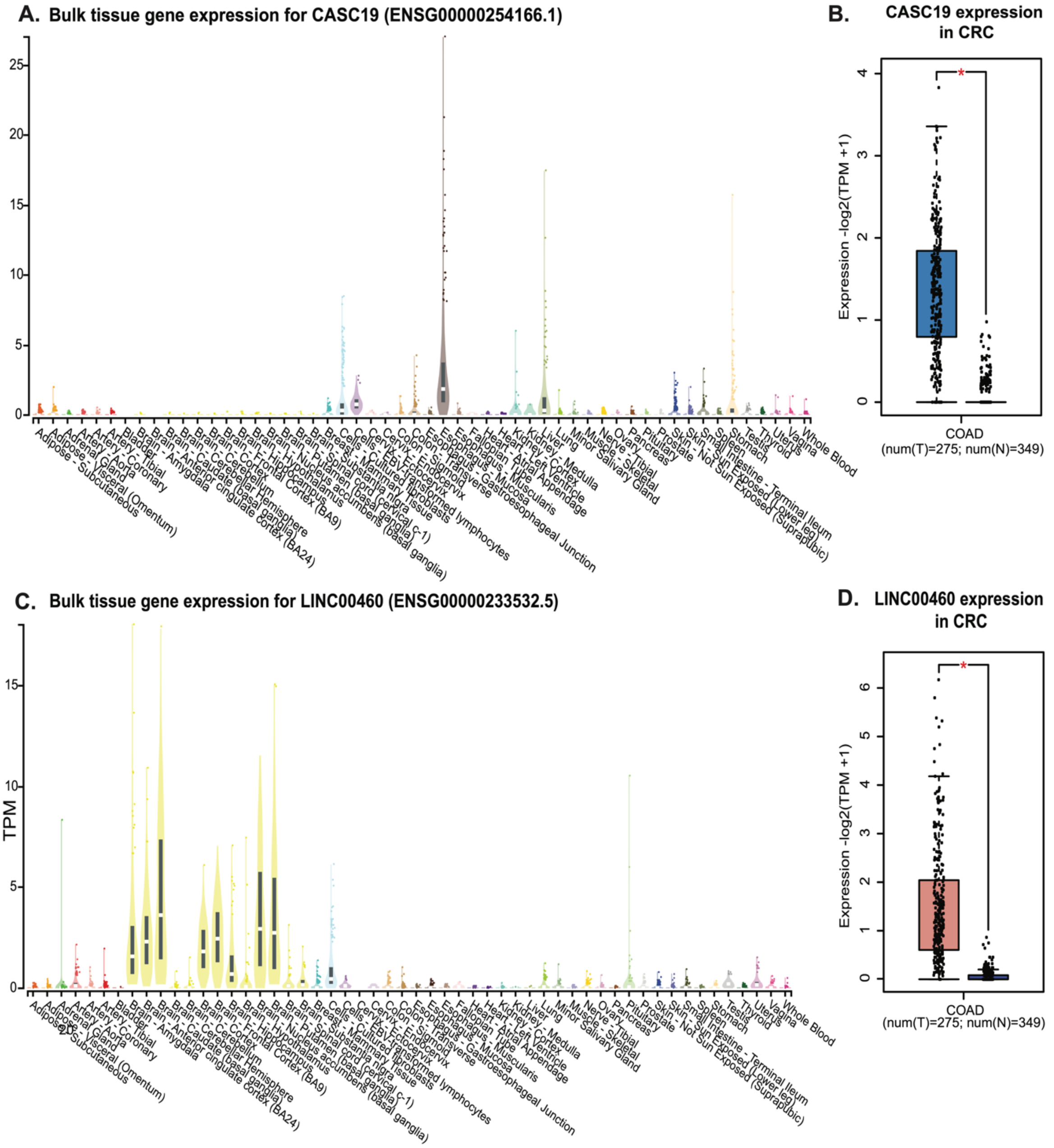


**Suppl. Figure 2. Bulk tissue gene expression of CASC19 and LINC00460 and their expression in CRC** A. and C. Bulk tissue gene expression for CASC19 (A.) and LINC00460 (C.). Data was obtained from GTEXportal. B. and D. The expression levels of CASC19 (B.) and LINC00460 (D.) between cancer and adjacent normal tissue in colon adenocarcinoma (COAD) based on The Cancer Genome Atlas (TCGA) database (T=tumour tissue, N=normal tissue).

**Suppl. Figure 3**

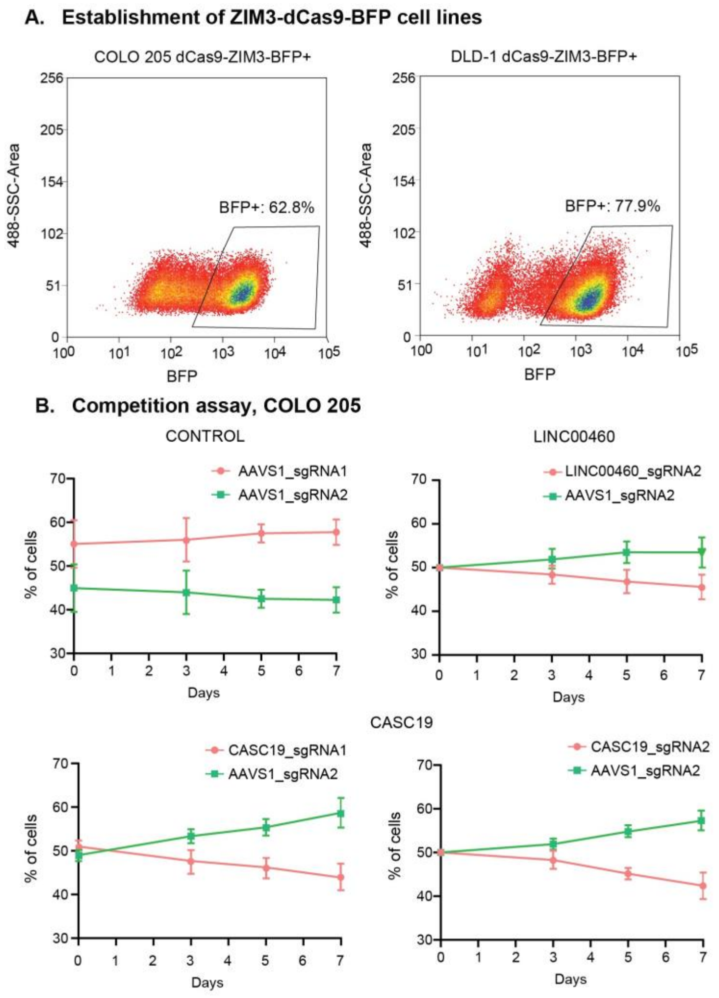


**Suppl. Figure 3. FACS mediated enrichment for ZIM3-KRAB-dCas9 expressing cells and competition assay in COLO 205 cell line.** A. Establishment of COLO 205 and DLD-1 cell lines stably expressing ZIM3-KRAB-dCas9 via FACS mediated enrichment for BFP expression. B. Competition assay used to assess the phenotypic implications of CASC19 and LINC00460 CRISPRi downregulation in COLO 205 cell line. Initially, cells expressing the mCherry marker and carrying various single-guide RNAs (sgRNAs) were combined with GFP-positive control cells in equal proportions. The red-to-green fluorescence ratio was measured at consecutive time intervals using flow cytometry. Data are plotted as mean ± SD (n=2).
